# Natural and laboratory mutations in *kuzbanian* are associated with heavy metal stress phenotypes in *Drosophila melanogaster*

**DOI:** 10.1101/070243

**Authors:** Hung Le Manh, Lain Guio, Miriam Merenciano, Quirze Rovira, Maite G. Barrón, Josefa González

## Abstract

Organisms must cope with altered environmental conditions such as high concentrations of heavy metals. Stress response to heavy metals is mediated by the metal-responsive transcription factor 1 (MTF-1), which is conserved from Drosophila to humans. MTF-1 binds to metal response elements (MREs) and changes the expression of target genes. *kuzbanian (kuz)*, a metalloendopeptidase that activates the evolutionary conserved *Notch* signaling pathway, has been identified as an MTF-1 target gene. We have previously identified a putatively adaptive transposable element in the *Drosophila melanogaster* genome, named *FBti0019170*, inserted in a *kuz* intron. In this work, we investigated whether laboratory-induced mutations in *kuz* are associated with zinc stress phenotypes. We found that both embryos and adult flies overexpressing *kuz* are more tolerant to zinc compared with wild-type flies. On the other hand, we found that the effect of *FBti0019170* on zinc stress tolerance depends on developmental stage and genetic background. Moreover, in the majority of the genetic backgrounds analyzed, *FBti0019170* has a deleterious effect in unpolluted environments in pre-adult stages. These results highlight the complexity of natural mutations and suggest that besides laboratory-induced mutations natural mutations need to be studied in order to accurately characterize gene function and evolution.

## INTRODUCTION

Heavy metals are non-degradable substances that are natural constituents of the Earth crust^1^. Some heavy metals, such as iron, copper, and zinc, are required at structural and catalytic sites in proteins and are thus vital for many biological processes such as transcription, respiration, and growth^1,2,3^. Indeed, heavy metal deficiencies are related to animal and human diseases such as neurodegenerative and cardiovascular disorders^2,4,5,6^. Although essential heavy metals are necessary for protein activity, when they are present at high concentrations they may bind to inappropriate sites in proteins interfering with their functions. Thus, under limiting conditions essential heavy metals have to be enriched while under excess conditions they have to be removed^7^.

Response to heavy metal stress is mediated by the metal-responsive transcription factor-1 (MTF-1). MTF-1 is conserved from insects to vertebrates and besides heavy metal stress it also mediates the response to oxidative stress and hypoxia^8,9,10,11,12^. MTF-1 binds to short DNA sequence motifs known as metal response elements (MREs) activating or repressing expression of target genes^13^. Functional MREs have been located in the promoter regions, downstream of the transcription start site, and in the introns of metal-responsive genes^7,13,14, 15^. In Drosophila, metallothioneins are the best characterized MTF-1 target genes. Metallothioneins have an extremely high affinity for heavy metal ions and play a role in both metal homeostasis and in defense against toxicity of heavy metals^14^.

Although their important role, a family knockout of all four Drosophila metallothioneins genes revealed that besides these proteins, other MTF-1 target genes must play a role in response to heavy metals and more specifically in zinc defense^16^. As expected according to these results, a genome-wide screen for MTF-1 target genes identified several candidate genes that respond to the presence of heavy metals in the environment^15^. One of these candidate genes, *kuzbanian* (*kuz)*, is a metalloendopeptidase that controls many biological processes during development and differentiation^17^. *kuz* belongs to the ADAM family of metalloendopeptidases that are zinc-dependent enzymes that have been shown to mediate stress response in mammals^18^.

In a previous work, we identified a putatively adaptive natural transposable element (TE), *FBti0019170*, inserted in an intron of *kuz* in *Drosophila melanogaster* natural populations^19^ (Fig. 1A). *FBti0019170* is a 4.7 kb non-LTR retrotransposon that belongs to the *F-element* family. *FBti0019170* is a strong candidate to play a role in the out-of-Africa adaptation: while most TEs are deleterious and thus present at low frequencies in populations, *FBti0019170* is present at high frequencies in North American populations and at low frequencies in African populations^19^. Additionally, the regions flanking this insertion showed signatures of a partial selective sweep. This suggests that *FBti0019170* has increased in frequency due to positive selection^19^. *FBti0019170* is located in the center of the sweep and we could not identify any other linked mutation further suggesting that the TE is the causal mutation. We have also already shown using allele-specific expression experiments that a *kuz* allele carrying *FBti0019170* insertion is overexpressed compared to a *kuz* allele without this insertion^19^.

**Figure 1.**
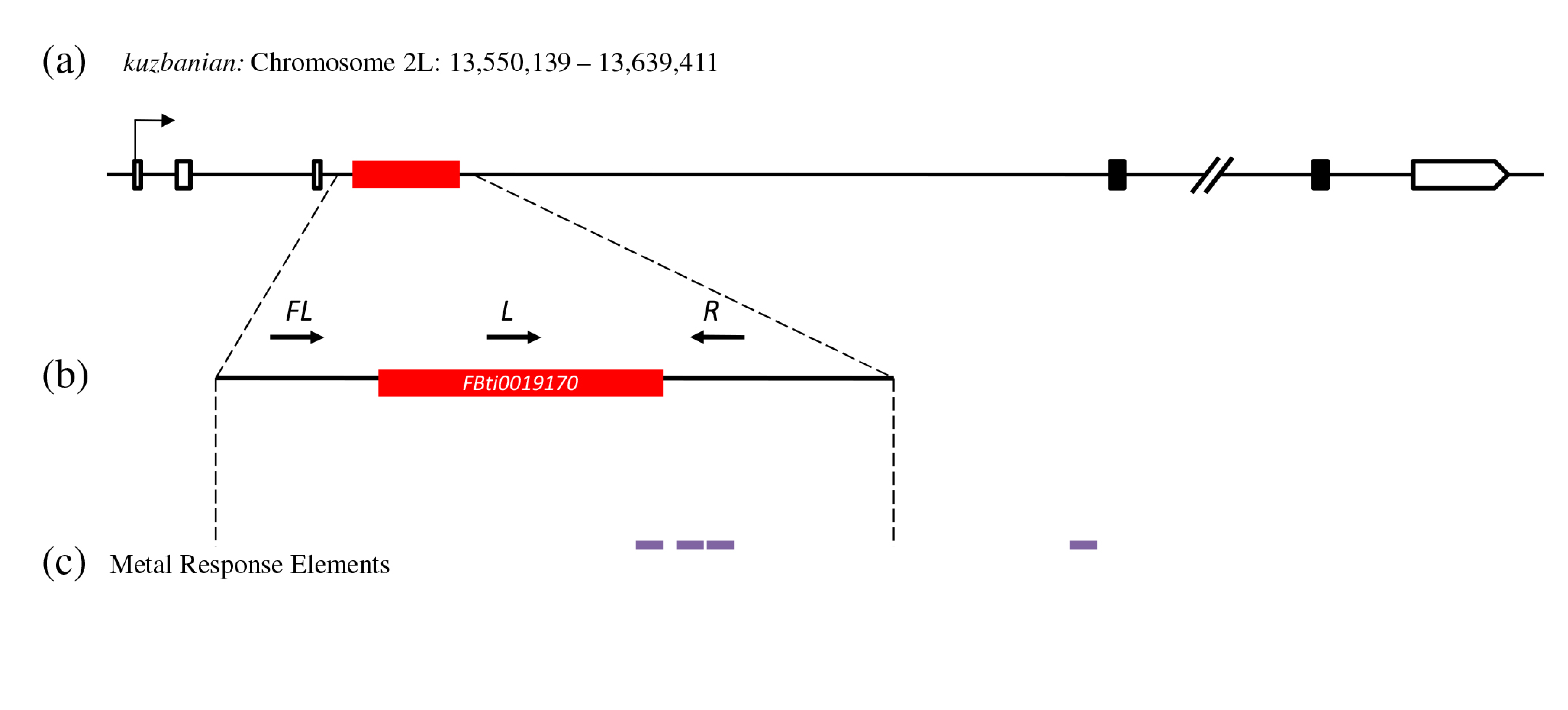
*FBti0019170* is a full-length *F-element*, 4,696 bp, inserted in the third intron of *kuzbanian.* (A) *kuzbanian (kuz)* gene region. White boxes represent UTRs, black boxes represent exons, black lines represent introns and intergenic regions, and the red box represents the *FBti0019170* insertion. (B) The region amplified represents *FBti0019170* insertion (2L: 13,560,515-13,565,210) and its flanking regions. Black arrows show the approximate localization of the three primers designed to check for the presence/absence of *FBti0019170*. (C) Predicted Metal Response Elements are represented in purple: one is located inside *FBti0019170* insertion and the other three in *kuz’s* third intron (see Table S2).

In this work, we investigated whether *kuz* is involved in heavy metal stress response, as has been previously suggested^15^, and whether *FBti0019170* has fitness consequences for flies that carry this natural insertion. We performed heavy metal stress tolerance assays using zinc chloride. High concentrations of zinc chloride are relevant for *D. melanogaster* natural populations because of its used in pesticides and fertilizers (www.epa.gov). We performed experiments with laboratory mutant flies and with flies collected in natural populations. Furthermore, because tolerance to environmental stress might differ between developmental stages, as has been already shown for lead, alcohol, and heat stress ^20,21,22,23,24^, we tested zinc stress tolerance in adult and pre-adult stages.

## RESULTS

### *kuz-overexpressing* flies are associated with increased zinc stress tolerance

To check whether *kuz* is involved in zinc stress response, we first determined the concentration of zinc that is required to kill 50% of wild-type flies (LD_50_). We tested 5 mM, 10 mM, and 20 mM and determined that 20 mM was the adequate dose for both male and female flies (Fig. S1A) (see Material and Methods). We then compared the survival rate of transgenic flies overexpressing *kuz, kuz-overexpressing* flies, with wild-type flies with a similar genetic background: *kuz-wildtype* flies (see Material and Methods). In nonstress conditions, that is, flies kept in standard fly food, we found no differences in the survival of *kuz-overexpressing* and *kuz-wildtype* flies (Fig. 2A). However, under zinc stress conditions, we found that *kuz-overexpressing* flies had higher survival than *kuz-wildtype* flies for both males and females (Fig. 2A and Table 1). To confirm these results, we performed a replicate of the experiment using flies from the same two strains a few generations later. We obtained the same results: both *kuz-overexpressing* male and female flies had higher survival than *kuz-wildtype* flies under zinc stress conditions (Fig. 2B and Table 1).

**Figure 2.**
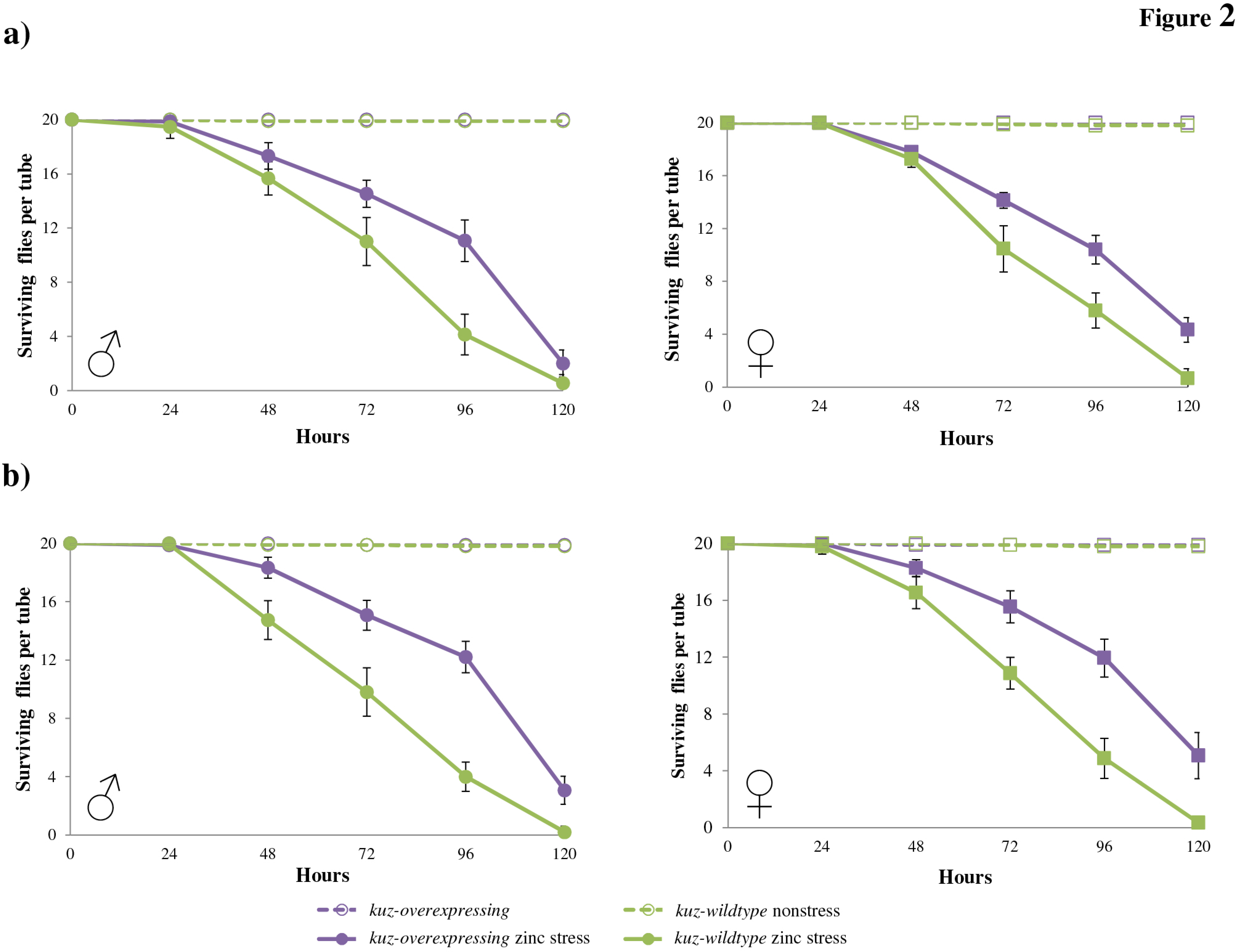
*kuz-overexpressing* flies are associated with zinc stress tolerance. Survival curves under nonstress (discontinuous lines) and under zinc stress (continuous lines) conditions are represented in purple for *kuz-overexpressing* flies, and in green for *kuz-wildtype* flies. The first replica (A) and the second replica (B) showed that *kuz-overexpressing* flies are more tolerant to zinc stress both in males and in female flies. Each data point in the survival curves represent the average survival for 15 tubes containing 20 flies each for zinc stress conditions and 10 tubes containing 20 flies each for nonstress conditions. In each data point, error bars represent the standard error of the mean (SEM).

**Table 1.**
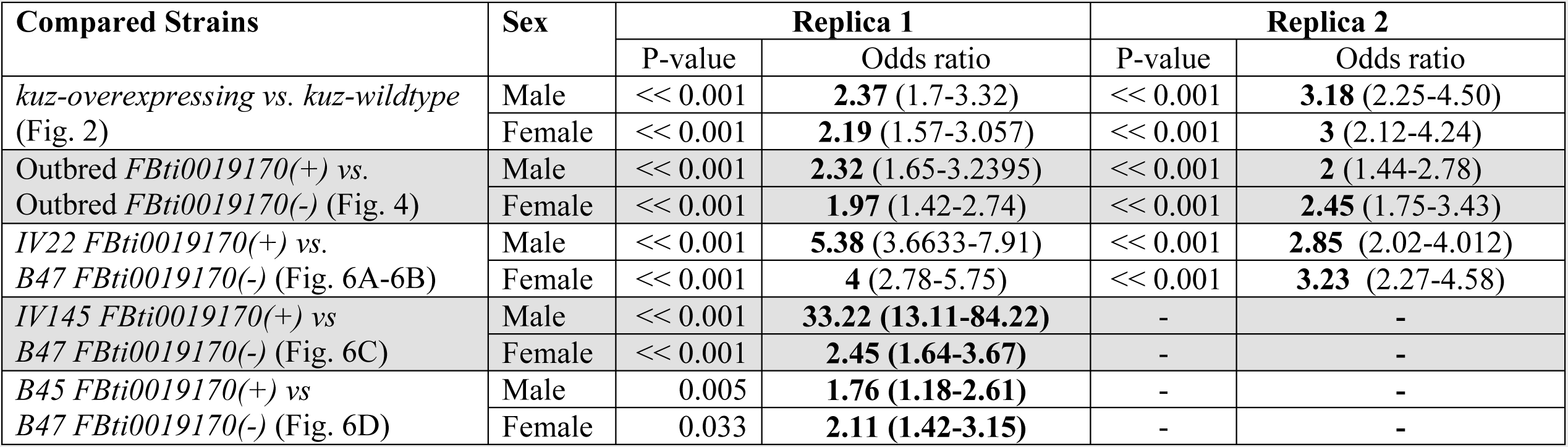
Log-rank analysis results and odds-ratio of heavy metal stress experiments performed with adult flies.

| Compared Strains | Sex | Replica 1 |  | Replica 2 |  |
| --- | --- | --- | --- | --- | --- |
|  |  | P-value | Odds ratio | P-value | Odds ratio |
| *kuz-overexpressing* vs. *kuz-wildtype*<br>(Fig. 2) | Male | << 0.001 | **2.37** (1.7-3.32) | << 0.001 | **3.18** (2.25-4.50) |
|  | Female | << 0.001 | **2.19** (1.57-3.057) | << 0.001 | **3** (2.12-4.24) |
| Outbred *FBti0019170*(+) vs.<br>Outbred *FBti0019170*(-) (Fig. 4) | Male | << 0.001 | **2.32** (1.65-3.2395) | << 0.001 | **2** (1.44-2.78) |
|  | Female | << 0.001 | **1.97** (1.42-2.74) | << 0.001 | **2.45** (1.75-3.43) |
| *IV22 FBti0019170*(+) vs.<br>*B47 FBti0019170*(-) (Fig. 6A-6B) | Male | << 0.001 | **5.38** (3.6633-7.91) | << 0.001 | **2.85** (2.02-4.012) |
|  | Female | << 0.001 | **4** (2.78-5.75) | << 0.001 | **3.23** (2.27-4.58) |
| *IV145 FBti0019170*(+) vs<br>*B47 FBti0019170*(-) (Fig. 6C) | Male | << 0.001 | **33.22** (**13.11-84.22**) | - | - |
|  | Female | << 0.001 | **2.45** (**1.64-3.67**) | - | - |
| *B45 FBti0019170*(+) vs<br>*B47 FBti0019170*(-) (Fig. 6D) | Male | 0.005 | **1.76** (**1.18-2.61**) | - | - |
|  | Female | 0.033 | **2.11** (**1.42-3.15**) | - | - |

Overall, we found that while there were no differences in survival between *kuz-overexpressing* and *kuz-wildtype* flies in nonstress conditions, *kuz-overexpressing* flies had higher survival than *kuz-wildtype* flies under zinc stress conditions (Fig.2 and Table 1). These results suggest that *kuz* could play a role in response to zinc stress.

### Egg to adult viability is higher in *kuz-overexpressing* flies in zinc stress conditions

As mentioned above, tolerance to environmental stress might differ between developmental stages. Thus, we also tested egg to adult viability under zinc stress conditions in *kuz-overexpressing* and *kuz-wildtype* flies. We first performed an LD_50_ and found that 10 mM zinc is the dose at which ~50% of the wild-type embryos do not emerge (Fig. S1B) (see Material and Methods).

We compared the egg to adult viability of *kuz-overexpressing* flies with *kuz-wildtype* flies in nonstress and in zinc stress conditions (Fig. 3). ANOVA analysis showed that the experimental condition (nonstress or zinc stress) and the interaction between experimental condition and genotype *(kuz-overexpressing* or *kuz-wildtype)* were significant (Fig. 3 and Table 2): *kuz-overexpressing* flies had higher egg to adult viability than *kuz-wildtype* flies in zinc stress conditions. This suggested that *kuz* could play a role in zinc stress response also in pre-adult developmental stages.

**Figure 3.**
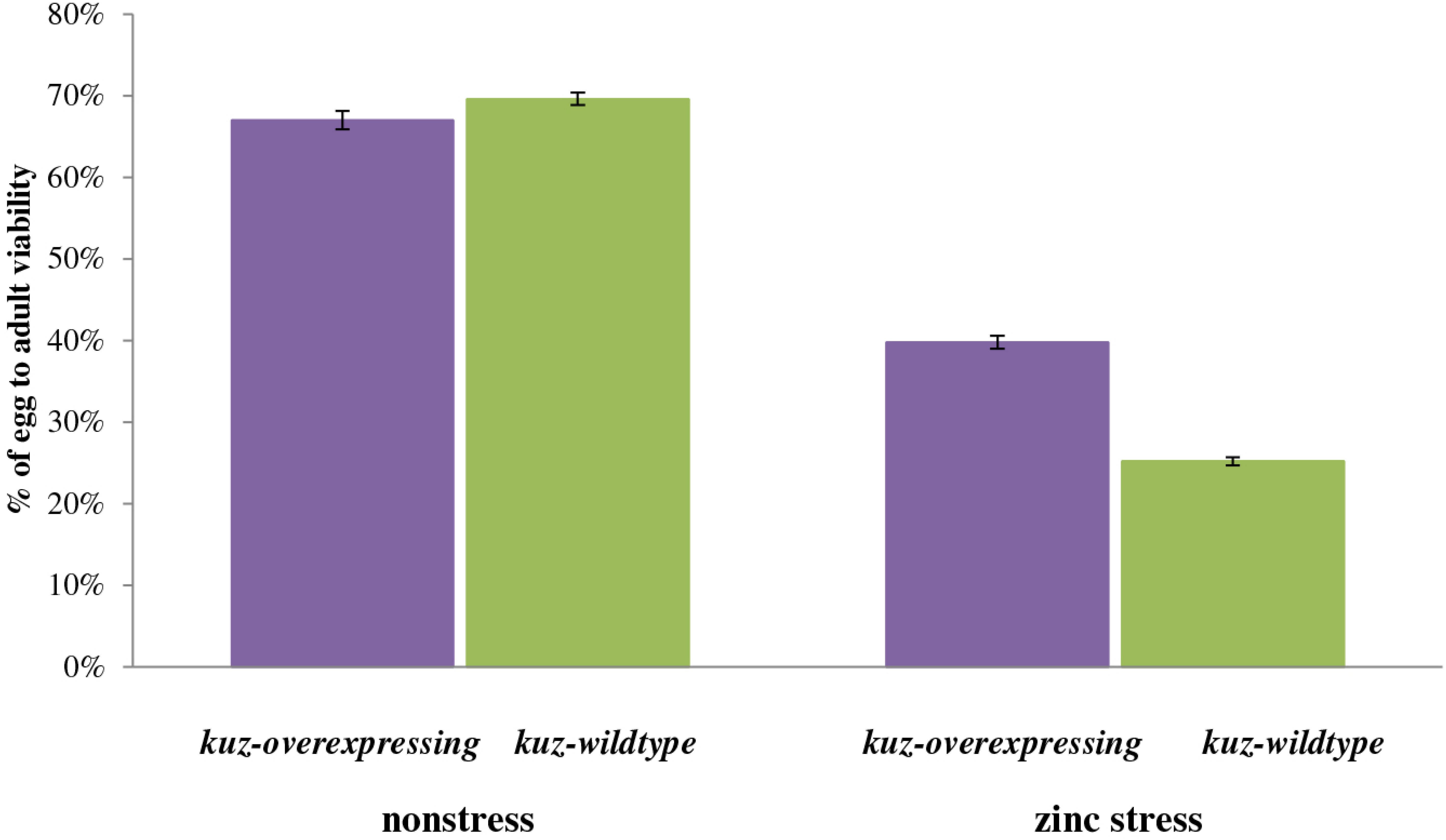
*kuz-overexpressing* embryos have a higher egg to adult viability under zinc stress. Each column represents the average of egg to adult viability. *kuz-overexpressing* flies are represented in purple and *kuz-wildtype* flies are represented in green. In each data point, error bars represent the standard error of the mean (SEM).

**Table 2.** Two-way ANOVA analyses of egg to adult viability experiments.

| Strains analyzed | Two-way ANOVA |  |  |  |  |  |
| --- | --- | --- | --- | --- | --- | --- |
|  | Experimental condition |  | Insertion genotype |  | Interaction between experimental condition & insertion genotype |  |
|  | P-value | Effect size | P-value | Effect size | P-value | Effect size |
| *kuz-overexpressing* vs *kuz-wildtype* (Fig. 3) | <<0.001 | 0.649 | 0.077 | - | 0.003 | 0.147 |
| Outbred *FBti0019170* (+) vs outbred *FBti0019170* (-) (Fig. 5) | <<0.001 | 0.688 | <<0.001 | 0.299 | 0.489 | - |
| *B47 FBti0019170* (-) vs *IV22 FBti0019170* (+) (Fig. 7A) | <<0.001 | 0.682 | 0.001 | 0.115 | <<0.001 | 0.138 |
| *B38 FBti0019170* (-) vs *B38 FBti0019170* (+) (Fig. 7B) | <<0.001 | 0.750 | 0.426 | - | <<0.001 | 0.146 |
| *IV52 FBti0019170* (-) vs *IV52 FBti0019170* (+) (Fig. 7C) | <<0.001 | 0.756 | 0.395 | - | 0.399 | - |

### *FBti0019170* is associated with increased zinc tolerance in flies from outbred populations

As mentioned above, *FBti0019170*, inserted in the third intron of *kuz*, shows signatures of a selective sweep suggesting that this natural insertion has increased in frequency due to positive selection. We thus investigate whether flies with *FBti0019170* were associated with increased tolerance to zinc stress. We analyzed both adult fly survival and egg to adult viability in nonstress and zinc stress conditions in natural strains with different genetic backgrounds: outbred strains and isofemale strains. Analyzing different genetic backgrounds is needed because the effect of mutations is often background dependent^25^.

We first constructed two outbred laboratory strains: one outbred strain homozygous for the presence of *FBti0019170* and one outbred strain homozygous for the absence of this insertion (see Material and Methods). We subjected these two outbred strains to zinc stress and we found that both male and female flies with *FBti0019170* had higher survival than flies without *FBti0019170* (Fig.4A, Table 1). The same results were obtained with the same outbred strains analyzed a few generations later (Fig.4B and Table 1). In both replicas, no differences in survival between flies with and without *FBti0019170* were found in nonstress conditions (Fig. 4). Overall, we found that *FBti0019170* is associated with increased zinc tolerance in adult flies.

**Figure 4.**
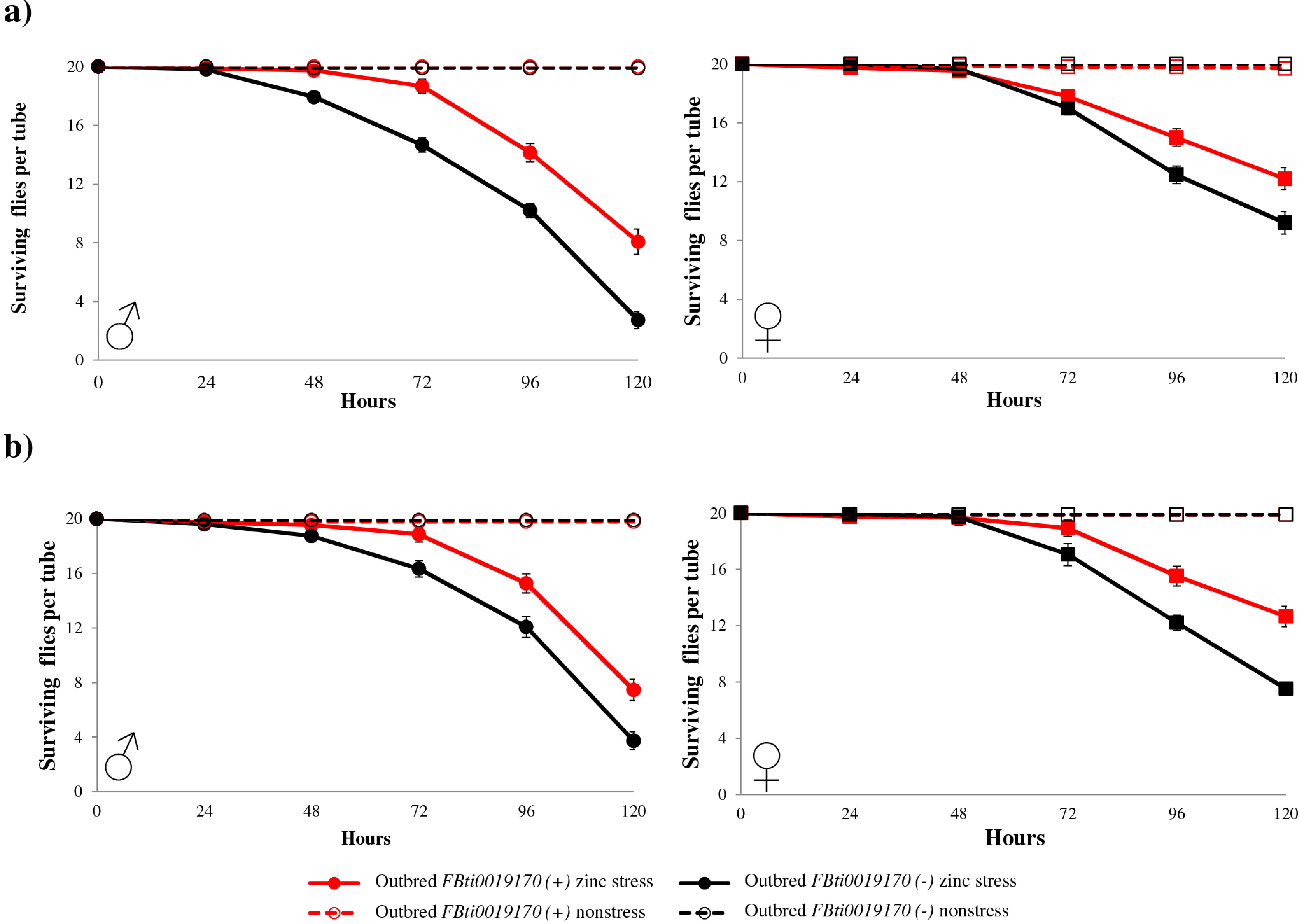
Outbred flies with *FBti0019170* insertion are associated with zinc stress tolerance. Survival curves under non-stress conditions (discontinuous lines), and under zinc stress (continuous lines) are represented in red for outbred flies with *FBti0019170* insertion, and in black for outbred flies without the insertion. The first (A) and the second replica (B) showed the same results for both males and females. Each data point in the survival curves represent the average survival for 15 tubes containing 20 flies each for zinc stress conditions and 10 tubes containing 20 flies each for nonstress conditions. In each data point, error bars represent the standard error of the mean (SEM).

### *FBti0019170* is associated with decreased egg to adult viability both in nonstress and in zinc stress conditions in outbred populations

We also tested whether embryos from the outbred strain with *FBti0019170* insertion were more tolerant to zinc stress compared with embryos from the outbred strain without the insertion. We performed two replicas of the experiment (Fig. 5). ANOVA analysis showed that the experimental condition (nonstress vs zinc stress) and the insertion genotype (presence vs absence of *FBti0019170*) were significant (Table 2). Both in nonstress and zinc stress conditions, flies with *FBti0019170* had a lower survival rate compared to flies without the insertion (Fig. 5). Thus, while *FBti0019170* is associated with higher adult survival in zinc stress conditions, it is also associated with lower egg to adult viability both in nonstress and in zinc stress conditions.

**Figure 5.**
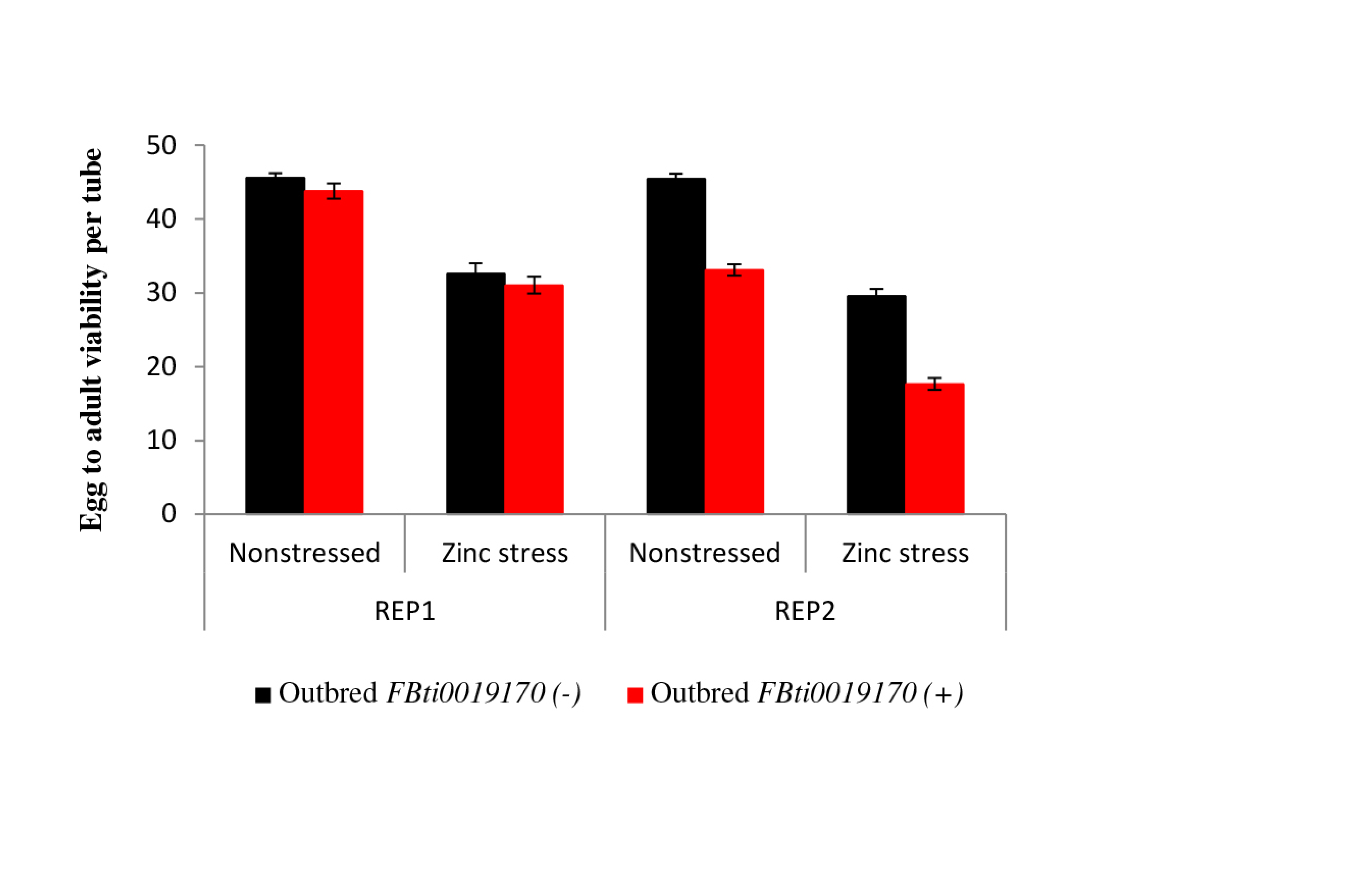
Egg to adult viability in outbred flies with *FBti0019170* is lower than in outbred flies without *FBti0019170* in nonstress and in zinc stress conditions. Each column represents the average of egg to adult viability of 15 vials for zinc stress conditions and 10 vials for nonstress conditions. Outbred flies without the insertion are represented in black and outbred flies with the insertion are represented in red.

### *FBti0019170* effect in adult survival in isofemale strains depends on the genetic background

We also performed adult survival experiments in nonstress and zinc stress conditions with different isofemale strains. Adult survival of *IV22, IV145*, and *B45* isofemale strains containing *FBti0019170* insertion was compared with the adult survival of *B47* isofemale strain without this insertion (see Material and Methods). No differences in survival between flies with and without *FBti0019170* insertion were observed in nonstress conditions. However, we found that *IV22* flies with *FBti0019170* had lower survival than *B47* flies without *FBti0019170* (Fig. 6A Table 1). We confirmed these results by performing a second replica a few generations later (Fig. 6B; Table 1). Similarly, *IV145* flies with *FBti0019170* also had lower survival than *B47* flies without *FBti0019170* (Fig. 6C Table 1). On the other hand, male flies of *B45* strain had higher survival than male flies of *B47* strain (Fig. 6D; Table 1) while *B45* female flies had higher survival than *B47* female flies at early time points while they had lower survival at later time points (Fig. 6D; Table 1).

**Figure 6.**
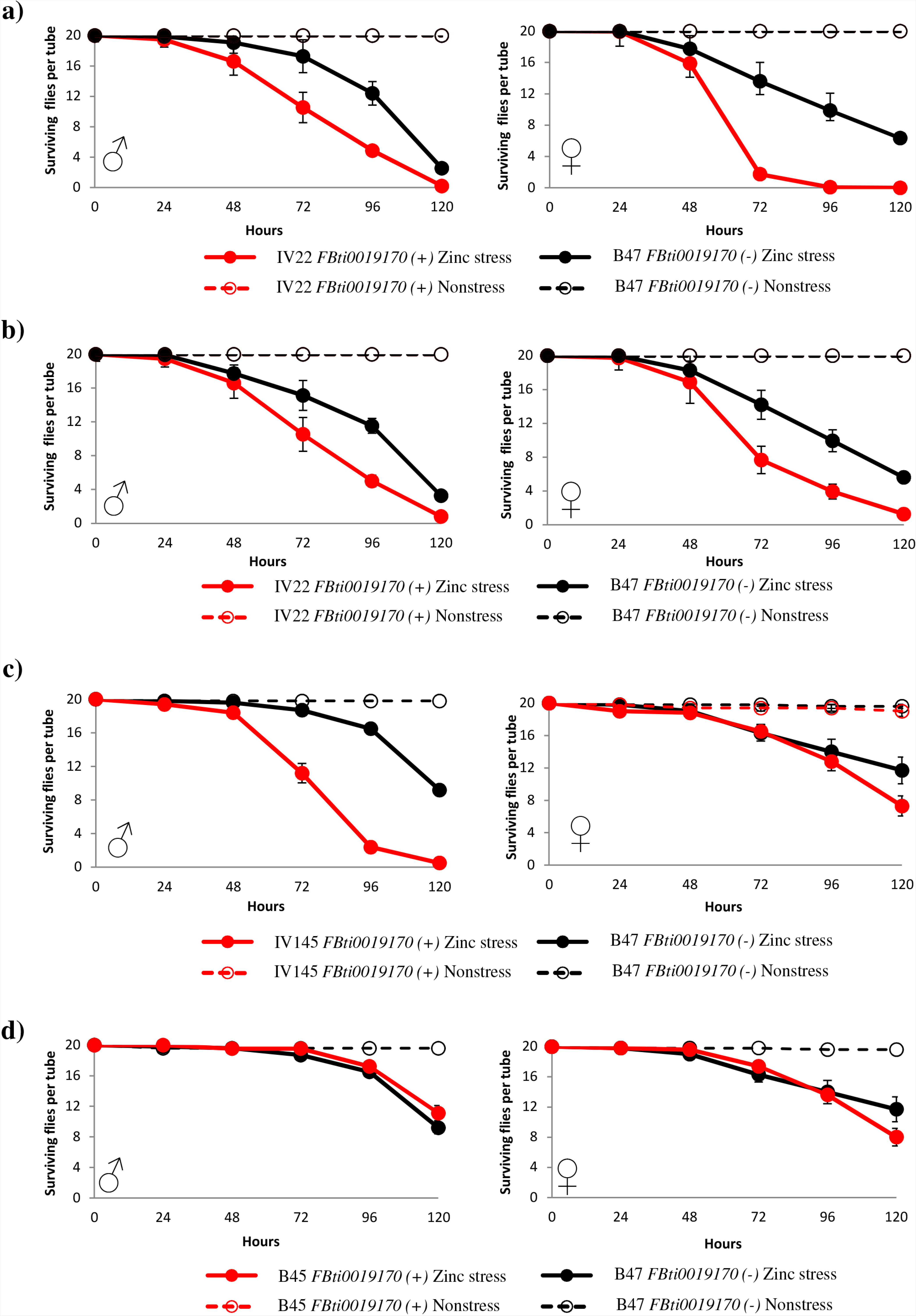
Isofemale strains with *FBti0019170* insertion are more sensitive or more tolerant to zinc stress depending on the genetic background. Survival curves under non-stress conditions (discontinuous lines) and under zinc stress (continuous lines) are represented in red for flies with *FBti0019170* insertion, and in black for flies without the insertion. (A) Survival curves for *IV22* vs *B47* (first replicate), (B) Survival curves for *IV22* vs *B47* (second replicate), (C) survival curves for *IV145* vs *B47*, and (D) survival curves for *B45* vs *B47.* Each data point in the survival curves represent the average survival for 15 tubes containing 20 flies each for zinc stress conditions and 10 tubes containing 20 flies each for nonstress conditions. Error bars represent the standard error of the mean (SEM) for each datapoint.

Therefore, in isofemale strains, the association between *FBti0019170* and heavy metal stress phenotypes depends on the background: while *IV22* and *IV145* isofemale strains with the insertion were more sensitive to zinc stress, *B45* were more tolerant to zinc stress in early time points compared to flies without this insertion.

### *FBti0019170* effect in egg to adult viability in isofemale strains also depends on the genetic background

We also checked egg to adult viability in the isofemale strain *IV22* that contains *FBti0019170* insertion and in *B47* strain without this insertion. Additionally, we identified two strains heterozygous for *FBti0019170, B38* and *IV52*, and we generated homozygous flies for the presence and homozygous flies for the absence of *FBti0019170* by brother-sister crosses (see Material and Methods). Note that flies with and without the insertion generated from heterozygous strains, have more similar genetic backgrounds compared with isofemale strains with and without the insertion.

For *IV22* and *B47*, we found that the experimental condition, the insertion genotype and the interaction between these two factors were significant (Figure 7A and Table 2). *IV22* flies that contain *FBti0019170* insertion had lower viability in nonstress conditions and higher viability in zinc stress conditions than flies without *FBti0019170.* For the *B38* flies with and without *FBti0019170* insertion, we found that the experimental condition, and the interaction between experimental condition and insertion genotype were significant (Fig. 7B; Table 2): in nonstress conditions flies with *FBti0019170* had lower viability while in zinc stress conditions had higher viability than flies without this insertion. Finally, for *IV52* strains with and without *FBti0019170*, only the experimental condition was significant (Fig. 7C; Table 2).

**Figure 7.**
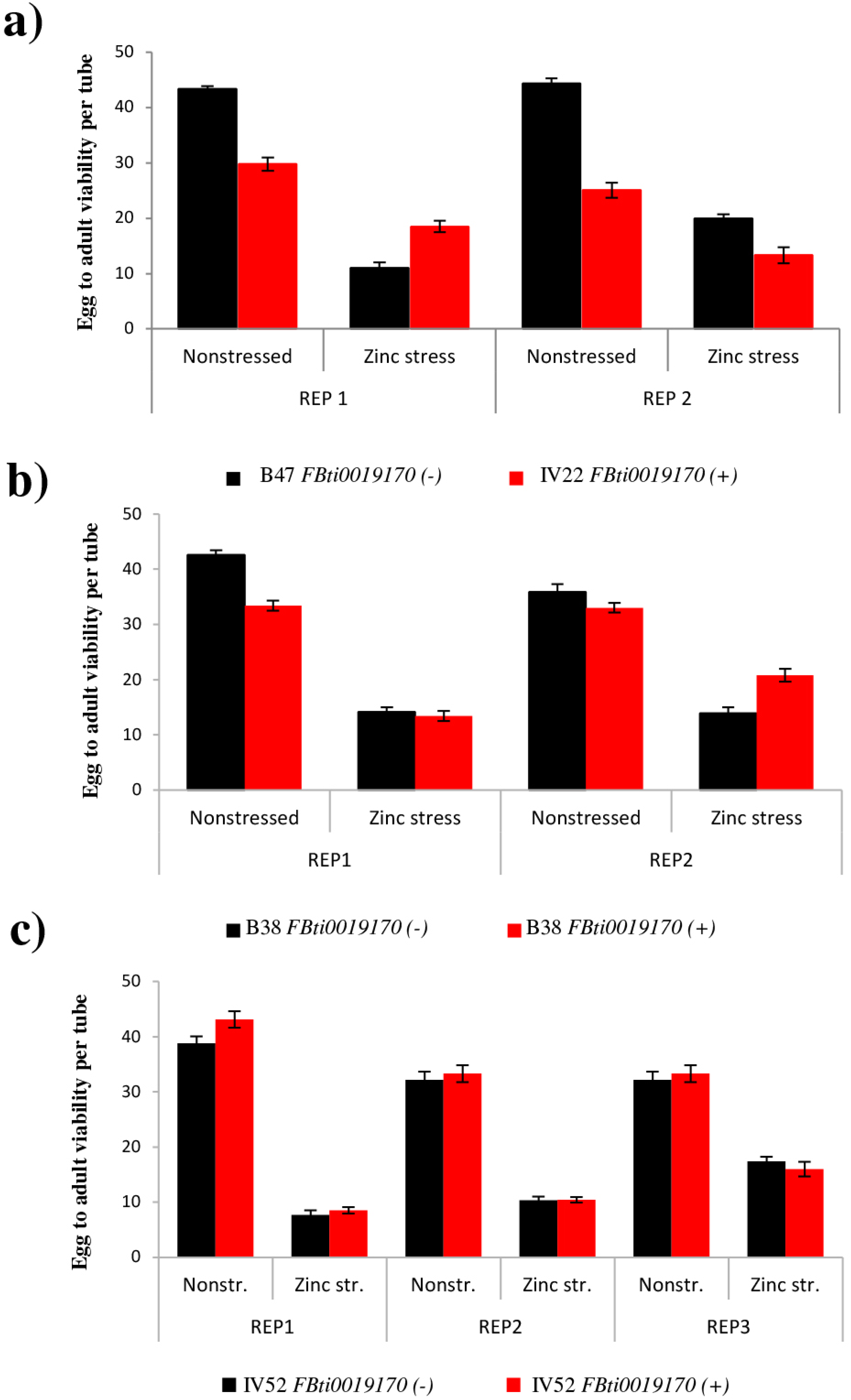
Egg to adult viability in isofemale strains with and without *FBti0019170* depends on the genetic background. Each column represents the average of egg to adult viability of 15 vials for zinc stress conditions and 10 vials for nonstress conditions. Strains without the insertion are represented in black and strains with the insertion are represented in red. (A) Egg to adult viability in *IV22* flies with *FBti0019170* and B47 flies without *FBti0019170*, (B) in *B38* flies with and without *FBti0019170* (C) and in *IV52* flies with and without *FBti0019170.*

Overall, we found that the effect of *FBti0019170* on egg to adult viability depends on the genetic background: in two of the three backgrounds analyzed the presence/absence of *FBti0019170* and/or the interaction between the genotype and the experimental condition was significant (Fig.7).

### *FBti0019170* could add a metal-responsive element to the intron of *kuzbanian*

As mentioned above, *kuz* was found to be a MTF-1 target gene, and we have shown that *kuz-overexpressing* flies were more zinc tolerant compared to *kuz-wildtype* flies. We have also found that flies with *FBti0019170* were associated with increased zinc tolerance in some genetic backgrounds. To shed light on the mechanisms underlying zinc stress response of laboratory-induced and natural *kuz* mutations, we investigated whether *kuz* has metal responsive elements (MREs) in its promoter region and whether *FBti0019170* is introducing any additional MRE. We could not detect any MRE in the *kuz* promoter. On the other hand, we found a high score MRE nearby the 3’ end of *FBti0019170* (Fig. 1C and supplementary Table S2). This prompted us to investigate whether there were other MREs in the *kuz* intron where *FBti0019170* is inserted, and we identified three additional MREs (Fig. 1C). Interestingly, two of these MREs are located only 462 bp downstream of the MRE introduced by *FBti0019170* while the third intronic MRE is located nearby the 3’ end of the intron (Fig. 1C and Table S2).

Overall, we found four MREs present in the *kuz* intron where *FBti0019170* is inserted. *FBti0019170* adds one of these four MREs. Because there is a correlation between the number of transcription factor binding sites and the increase in the level of expression of nearby genes^26^, these results suggest that *FBti0019170* could affect the expression of *kuz* and thus could play a role in zinc stress response.

### Flies homozygous for the presence and for the absence of *FBti0019170* insertion do not show differences in the level of *kuz* expression

We checked whether outbred flies homozygous for the presence of *FBti0019170* showed different levels of *kuz* expression compared to outbred flies without this insertion. We performed qRT-PCR experiments both in nonstress and in zinc stress conditions for both male and female flies. No differences in the level of expression of *kuz* between flies with and without *FBti0019170* were found in nonstress or zinc stress conditions for males or for females (Fig. 8).

**Figure 8.**
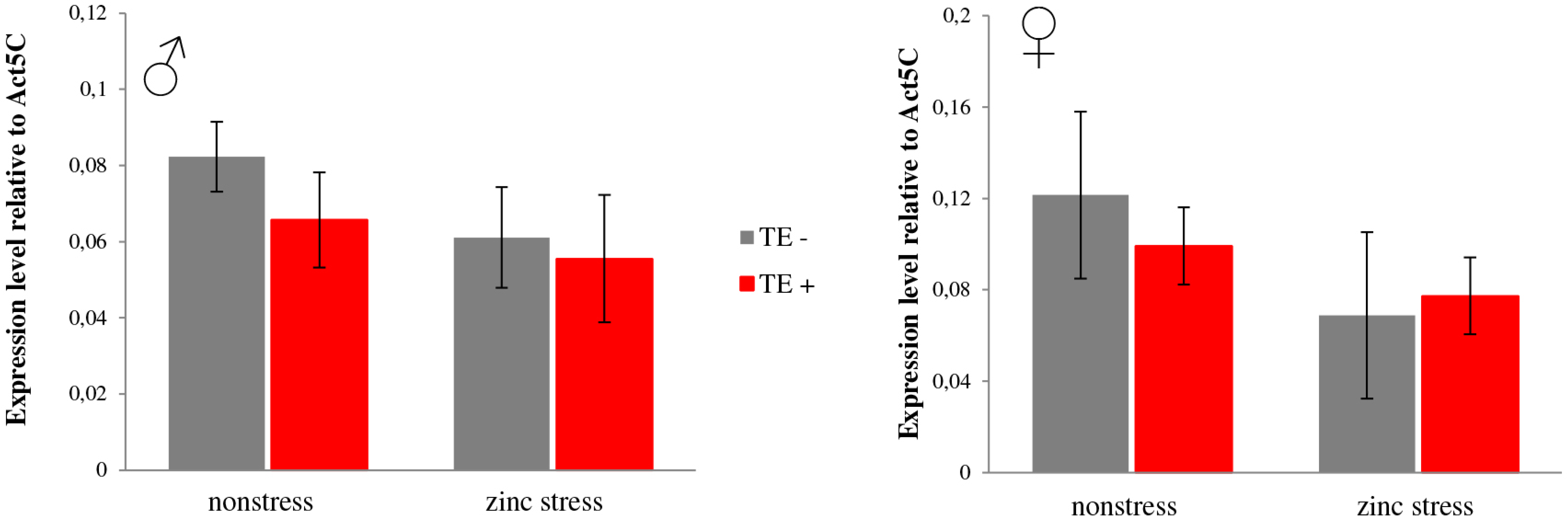
Flies homozygous for the presence and for the absence of *FBti0019170* showed no differences in *kuz* expression. Normalized expression level relative to *Act5C* of *kuz* in nonstress and zinc stress conditions in male and female flies. Flies without *FBti0019170* insertion are represented in grey and flies with *FBti0019170* are represented in red. Error bars represent the SEM of three biological replicas.

## DISCUSSION

In this work, we showed that a laboratory-induced mutation in *kuz* is associated with tolerance to zinc stress both in adult (Fig. 2) and embryo stages (Fig. 3). These results are consistent with a role of *kuz* in heavy metal stress response, as it has been previously suggested by experiments performed with MTF-1 mutant flies that identified *kuz* as a candidate heavy metal-responsive gene^15^. *kuz, a* metalloprotease that belongs to the ADAM family, is a component of the Notch signaling pathway that plays a role in axon guidance in the developing central nervous system^17,27-30^. ADAM metalloproteases in mammals, and more specifically *kuz* ortholog ADAM10, ADAM17, and to a lesser extend ADAM9, also regulate epidermal growth factor receptor (EGFR) activation in response to a variety of stress agents^18^. Stress-induced EGFR activation leads to the activation of mitogen-activated protein kinase (MAPKs) signaling that trigger transcriptional regulation of a variety of stress-response genes^1^. Thus, both *kuz* and its ortholog gene ADAM10 could be involved in response to stress. Indeed, the organomercurial compound p-aminophenylmercuric acetate (APMA) have been reported to upregulate both *kuz* and *ADAM10* protease activity^31,32^ and methylmercury has been suggested to activate ADAM proteases in Drosophila^33^.

Our previous results showing that *FBti0019170*, which is inserted in the third intron of *kuz*, has most probably increased in frequency due to positive selection, prompted us to investigate whether flies with this insertion have increased zinc tolerance. To test this hypothesis, we generated an outbred population and analyzed five isofemale strains established from two different natural populations. We found that the effect of *FBti0019170* on egg to adult viability and on adult survival under zinc stress conditions depended on the genetic background analyzed (Figure 4-7). In four of the six backgrounds analyzed, the presence of *FBti0019170* insertion was associated with higher adult survival or increased egg to adult viability under zinc stress conditions. In the other two backgrounds, *FBti0019170* was not significantly associated with zinc stress phenotypes or *FBti0019170* was associated with increased sensitivity to zinc stress (Fig. 4-7). These results are consistent with previous experimental data showing that the mutational effects in one genetic background are often enhanced or suppressed in other backgrounds^34,35^. For example, different mutations in the same gene have been associated with both increased and decreased life span in *D. melanogaster^36^.* Different effects of a mutation, as the ones described in this work, are most likely explained by epistatic interactions in the different genetic backgrounds^25^.

Our results also showed that *FBti0019170* was associated with lower egg to adult viability in nonstress conditions in three of the four backgrounds analyzed (Fig. 5 and 7). Between-environments trade-offs have been reported for cadmium resistance in *D. melanogaster*^37^ as well as for other environmental stress conditions such as oxidative stress^38^. Two mechanisms have been proposed to explain the fitness costs of heavy-metal resistance in unpolluted environments: the activation of detoxification enzymes might use resources that are then unavailable for other fitness traits, and/or resistant flies might be less efficient at metal uptake or utilization, which would lead to micronutrient deficiencies^37^. In the case of *FBti0019170*, the deleterious effect of the mutation was found in egg to adult viability while no cost of selection was found in adult stages. As mentioned above, *kuz* plays a role in development and differentiation^17,27-30^. Thus, it could be that the cost of selection of *FBti0019170* is related to the role of *kuz* during development.

Consistent with the activation of *kuz* by zinc, we found *in silico* evidence for three MREs located in the *kuz* intron where the candidate adaptive TE *FBti0019170* is inserted (Fig. 1C). Indeed, *FBti0019170* insertion adds another MRE 462 bp upstream of the three intronic MREs (Fig. 1C). We thus check the expression of *kuz in* flies homozygous for the presence and for the absence of *FBti0019170* using qRT-PCR. We did not find differences in *kuz* expression in nonstress or in zinc stress conditions (Fig. 7). However, we have previously shown, using allele-specific expression, that an allele carrying *FBti0019170* insertion is overexpressed compared to an allele that does not carry this insertion^19^. Because allele-specific expression is performed in F_1_ hybrids in which the two alleles share the same cellular environment, the expression differences between the two alleles must be due to cis-regulatory differences^39^. *FBti0019170* is thus a strong candidate to be responsible for the observed differences in *kuz* expression level^19^. The lack of differential expression in homozygous flies could be partly explained by the higher sensitivity of allele-specific expression experiments compared to qRT-PCR^40^. Besides, it could be that *FBti0019170* effect on *kuz* expression is overdominant as has been described for a few genes involved in temperature stress response^41^. Further experiments are needed in order to understand the molecular mechanism underpinning *FBti0019170* insertion effects.

As with other quantitative traits, including starvation stress and olfactory behavior, we have found that mutations in a gene with well-characterized roles in development affect tolerance to zinc stress^34^. Our results showed that while *kuz* laboratory-induced mutations are consistently associated with increased tolerance to heavy metal stress in embryo and adult stages, natural mutations have more complex fitness effects that depend on the developmental stage and the genetic background. Furthermore, while no cost of selection was associated with the laboratory-induced mutation, we found that *FBti0019170* is associated with decreased egg to adult viability in unpolluted environments. Different fitness effects of laboratory-induced and natural mutations have previously been described suggesting that the analysis of natural mutations is needed in order to accurately characterize gene function and evolution^42,43^.

## METHODS

### Genotyping flies for presence/absence of *FBti0019170.*

To check the insertion genotype, that is, whether different fly stocks were homozygous for the presence, homozygous for the absence, or heterozygous for *FBti0019170* insertion, we performed PCR with two pairs of primers (^19^. Primer pair *Left* (L) and *Right* (R) were designed to check for the presence of *FBti0019170* (Fig.1B). The *Left* primer (TTCGGAGTGAAAACATCCAAAGA) binds to *FBti0019170* while the *Right* primer (TTGAATATTGTGTCGATTGCGTG) binds to the downstream sequence flanking the insertion (Fig.1B). This primer pair only gives a PCR band when *FBti0019170* is present. On the other hand, primer pair *Flanking* and *Right* was designed to check for the absence of *FBti0019170*. The *Flanking* (FL) primer (GACGAATTCATAAATTGGCGGTT) binds to the upstream sequence flanking the insertion (Fig.1B). This primer pair only gives a PCR band when *FBti0019170* is absent. If only the *Left-Right* primer pair gives a PCR band, the strain is homozygous for the presence of *FBti0019170.* If only the *Flanking-Right* primer pair gives a PCR band, the strain is homozygous for the absence of *FBti0019170.* Finally, if both primers give PCR bands, the strain is heterozygous for *FBti0019170* insertion^44^. 48 different isofemale strains collected in Stockholm (Sweden, “B” strains) and 15 isofemale strains collected in Bari (Italy, “IV” strains), available in our laboratory, have been tested by PCR to check for the presence/absence of *FBti0019170* natural insertion (Table S1).

### Fly strains

#### Laboratory mutant strains

We used transgenic flies that carry a full copy of *kuz* coding region under the control of a *UAS* promoter^45^ (Bloomington stock # 5816). To activate the expression of *kuz*, we crossed the flies with transgenic flies that carry the *GAL4* gene under the control of *Act5C* promoter (Bloomington stock # 4414) and we kept the crosses at 25°C. A total of 200 virgin female of *kuz* mutant flies were crossed with 200 male of *Act5C-GAL4* flies. F_1_ flies carrying *UAS-kuz* and *Act5C-GAL4*, and thus overexpressing *kuz*, have wild-type wings *(kuz-overexpressing* flies) while F_1_ flies that do not have the *Act5C-GAL4* chromosome and thus do not over-express *kuz* have *Curly* wings. A stock with a w[*] genetic background, as the *kuz* transgenic flies background, was used as the baseline for the experiment *(kuz-wildtype* flies; Bloomington stock # 7087).

#### Outbred populations

To create an outbred population with *FBti0019170* insertion and an outbred population without the insertion, a total of 10 isofemale strains were selected: five strains homozygous for the presence of the element *(B7, B45, IV33, IV49, IV50)* and five strains homozygous for the absence *(B2, B4, B8, B15, B18)* (Table S1). We collected 10 virgin females and 10 males from each one of these strains. We did two crosses by mixing males and females with the TE to create an outbred *FBti0019170 (+)* strain, and males and females without the TE to create an outbred *FBti0019170 (-)* strain. We kept the two populations for at least seven generations before performing any phenotypic experiments.

#### Isofemale Strains

We selected three strains in which *FBti0019170* was present *(IV22, IV145* and *B45*) and one strain in which *FBti0019170* was absent *(B47)* to perform phenotypic experiments (Table S1). Isofemale flies heterozygous for *FBti0019170* insertion were also selected to create homozygous flies for the presence and homozygous flies for the absence of *FBti0019170* (see below).

#### Heterozygous strains

We first identified two isofemale strains *(B38* and *IV52)* that were heterozygous for *FBti0019170* insertion. We then collected 10 to 25 virgin females from each strain and crossed them individually with males from the same strain. F_1_ progeny of all the crosses were checked for the presence/absence of the *FBti0019170* insertion by PCR. Brother-sister crosses were performed until we obtained flies that were homozygous for the presence of *FBti0019170* and flies that were homozygous for the absence of *FBti0019170.* Those flies were amplified for several generations in order to obtain enough quantity of flies to perform the experiments.

### Zinc stress experiments

We used zinc chloride (ZnCl_2_) as a heavy metal stress agent (Sigma-Aldrich catalog # Z0152). We have performed zinc stress experiments in two different life stages: adult flies and embryos.

#### Adult flies

To determine the Lethal Dose_50_ (LD_50_) for the adult flies experiments, we tested three different zinc concentrations: 5 mM, 10 mM and 20 mM. The experiments allowed us to identify the ZnCl_2_ concentration at which about 50% of the adult flies die. ZnCl_2_ was dissolved in water and added to the fly food to the desired final concentration. Standard fly food was used for the nonstress conditions. We used the outbred population without *FBti0019170* to establish the LD50. We analyzed 10 vials for each concentration and sex with 20 five to seven day-old flies each.

For the zinc stress tolerance experiments with adult flies from natural populations, a total of 100 vials with 20 flies each were used including 40 vials for nonstress conditions (10 vials per sex and per strain) and 60 vials for zinc stress condition (15 vials per sex and per strain). We used five to seven day-old flies.

For the zinc stress experiments performed with *kuz-overexpressing* flies, we used 40 vials for the nonstress condition and 60 vials for the zinc stress condition. 10 vials per condition were used to perform the experiments for the *kuz-wildtype* flies. We used five to seven day-old flies.

#### Embryos

We determined the LD50 using embryos from an isofemale strain without *FBti0019170* insertion (B47). These experiments allowed us to identify the ZnCl_2_ concentration at which about 50% of the embryos do not emerge. We first tested three different ZnCl_2_ concentrations: 1.25 mM, 2.5 mM and 5 mM. The same strain and protocol but different concentrations of ZnCl_2_ were tested in a second LD_50_ experiment: 5 mM, 10mM, 20 mM. In both cases, standard fly food was used for the nonstress conditions. Five day-old isofemale flies without *FBti0019170* insertion were kept in chambers with agar and apple juice plates to lay eggs during 4 hours. 10 vials with 50 embryos each were analyzed for each of the three ZnCl_2_ concentrations and for the nonstress condition.

Once the LD_50_ was determined, we performed zinc stress experiments using 50 embryos per vial. For the *kuz-wildtype* strain, we analyzed 10 vials for nonstress and 10 vials for zinc stress conditions. For the *Kuz-overexpressing* strains, we analyzed 20 vials for nonstress and 20 vials for zinc stress conditions. For natural strains, we used 15 vials for the zinc stress condition and 10 vials for the nonstress condition.

### *In silico* prediction of MTF-1 binding sites

We use TFBSTools software^46^ to predict binding sites of MTF-1 in the *kuz* promoter region and in the *kuz* intron where *FBti0019170* is inserted. Position weight matrices for MTF-1 transcription factor were obtained from JASPAR database^47^. The PB0044.1 and PB00148.1 matrices were used. Although the default score threshold in TFBSTools is 0.75, we were conservative and we only considered significant those hits with a score threshold ≥ 0.95. We used the release 6.02 of the *D. melanogaster* genome available at http://flybase.org.

### qRT-PCR Expression analysis

We quantified the expression of *kuz* in nonstress and stress conditions induced by zinc. Five day-old outbred flies (30 females and 50 males) were separated by sex and transferred to standard fly food as well as food containing 20 mM zinc for 48 hours before freezing them in liquid nitrogen. We did three biological replicas with flies from three different generations for each sex and condition. We purified total RNA using Trizol reagent and we synthesized cDNA using μg of RNA after treatment with DNase. We then used the cDNA for quantitative PCR analysis using *Act5C* as a housekeeping gene. The primers used were as follows: *kuz_left primer:* CACCGAGCATCGCAACATAC, *kuz_right primer:* GAATTGCGACAGGCCGAATC, *Act5C_left primer:* ATGTCACGGACGATTTCACG, and *Act5C_right primer:* GCGCCCTTACTCTTTCACCA.

Results were analyzed using the dCT method and following the recommendations of the MIQE guideline^48^.

### Statistical analysis

#### Log-rank test

The number of surviving flies for both nonstress and stress conditions were counted every 24 hours for five consecutive days. We used Kaplan-Meier to estimate the survival functions and performed a log-rank test to compare the functions between flies with and without the insertion using the SPSS software.

The odds-ratio (O.R.) was calculated as: (number of tolerant flies alive/number of tolerant flies dead)/ (number of sensitive flies alive/number of sensitive flies dead). The upper and lower 95% O.R. confidence interval was calculated as: e ^ [ln OR ± 1.96 √ (1/ number of tolerant flies alive + 1/ number of tolerant flies dead + 1/ number of sensitive flies alive + 1/ number of sensitive flies dead)]. We used the 95% confidence interval as a proxy for the presence of statistical significance when it does not overlap the null value, that is, O.R. = 1.

#### Two-way ANOVA analyses

The number of flies emerging from the experiments performed with embryos was transformed to a uniform distribution using the rank transformation. SPSS v21 was used to perform the ANOVA analyses. Two different variables were considered for the ANOVA analyses: the genotype *(kuz-overexpressing/ kuz-wildtype* or *FBti0019170* present/ *FBti0019170* absent), and the experimental condition (nonstress and zinc stress). The replicate effect was also considered. As a measure of the effect size, we estimated partial eta-squared values (0.01 small effect, 0.06 medium effect, and 0.14 large effect).

## ACKNOWLEDGEMENTS

We thank members of the González lab for their comments on the manuscript. H.L.M. was a VAST-CSIC fellow, L.G. was a FI/DGR fellow (2012FI-B-00676) and J.G. is a Ramón y Cajal fellow (RYC-2010-07306). This work was supported by grants from the European Community’s Seven Framework Programme (FP7-PEOPLE-2011-CIG-293860), from the Spanish Government (BFU2011-24397 and BFU2014-57779-P), and from the Generalitat de Catalunya (2014 SGR 201).

## AUTHOR CONTRIBUTIONS

H.L.M. performed research, analyzed data, and draft the manuscript. L.G., M.M., Q.R and M.G.B, performed research and analyzed data. J.G. designed research, analyzed data, and wrote the manuscript.

